# The effects of temporal smoothing and interpolation on EEG microstate parameters, syntax, and our view of resting-state brain dynamics

**DOI:** 10.64898/2026.09.30.755764

**Authors:** Frederic von Wegner, Gesine Hermann, Inken Tödt, Inga Karin Todtenhaupt, Helmut Laufs

## Abstract

Microstate analysis studies dynamic brain activity through a discrete representation of electroencephalographic (EEG) signals and their topography. While the body of literature describing findings in diverse experimental and clinical settings is growing rapidly, the effects of several core components of the microstate algorithm are still insufficiently understood. This study systematically investigates how two common temporal smoothing algorithms, parametric smoothing and GFP peak interpolation, influence a wide range of microstate sequence parameters, and how they shape the integrated picture of brain dynamics inferred from these metrics.

We study eyes-closed resting-state EEG recordings from N = 201 healthy subjects from a public EEG database. For the elementary temporal metrics, we find that widening smoothing windows markedly increase microstate duration, reduce occurrence rates, but leave coverage mostly unaffected. In terms of simple syntax properties, smoothing windows above 30 ms yield an inconsistent picture with a memoryless, first-order Markov structure at short time scales, but a positive Hurst phenomenon suggestive of long-range correlations at long time scales. Evidence of asymmetry and non-equilibrium dynamics are suppressed by smoothing. At intermediate time scales, microstate periodicity was only observed up to a window size of 28 ms. Smoothing had a strong impact on the complexity metrics entropy rate and excess entropy, decreasing the former and increasing the latter. For most metrics, the effects of GFP peak interpolation were comparable to those of smoothing windows in the range between 28 ms and 44 ms.

In conclusion, elementary and higher-order microstate parameters must be interpreted in the context of preprocessing, as many parameters change continuously with smoothing intensity. This limits the comparability of microstate parameters obtained with different preprocessing settings. Strong smoothing with a window size above 30 ms or GFP peak interpolation yields a fundamentally different picture of resting state brain dynamics, i.e. that of a memoryless, aperiodic, time-reversible first-order Markov chain. Since insights from other neurophysiological modalities suggest that time series memory and time-irreversibility are markers of normal brain activity, our recommendation is to avoid intense smoothing to minimize alterations of the initial sequences.

## Introduction

Electroencephalographic (EEG) microstate analysis has become a well-established approach for studying the rapid temporal dynamics of large-scale brain electrical activity (Michel and Koenig, 2018). Over the past decade, the use of microstates has expanded rapidly, with a growing body of literature documenting alterations in microstate parameters across a wide range of physiological conditions and clinical populations. Several recent reviews and metaanalyses have synthesized findings across this expanding literature (Michel et al, 2024). To give some examples, Michel and Koenig (2018) survey the history and current state-of-the-art of microstate research, and compare alterations in temporal microstate parameters across 19 neuropsychiatric resting-state studies, while event-related microstate studies across different psychiatric disorders are reviewed in Perrottelli et al (2025). Another study conducted a meta-analysis of microstate alterations in mood and anxiety disorders (Chivu et al, 2024), Bréchet and Michel (2022) reviewed altered states of consciousness, and Tarailis et al (2023) summarize our current understanding of the functional role of different microstate classes. Overall, this expansion reflects increasing recognition that microstates capture fundamental building blocks of time-varying neural activity and may serve as potential biomarkers for psychopathology and cognition.

Yet, at the same time that our understanding of how different physiological and pathological conditions affect microstate parameters is growing, our understanding of how certain parameter settings at the core of the microstate processing pipeline shape microstate parameters is very limited. In particular, the effects of microstate sequence smoothing have not been studied systematically (Michel and Koenig, 2018). Originally motivated by the need to reduce noise in pointwise label assignments, the aim of smoothing is to redefine the microstate boundaries and is therefore expected to significantly influence microstate metrics (Pascual-Marqui et al, 1995; Koenig et al, 1999).

Two smoothing approaches have been particularly influential. The first is parametric temporal smoothing, introduced as part of the canonical microstate segmentation and backfitting procedure in Pascual-Marqui et al (1995). The algorithm achieves smoothing iteratively, minimizing a cost function that contains a penalization term against rapid microstate switches. The algorithm contains two free parameters, the window half-size and a strength parameter. The second, GFP peak interpolation, is non-parametric and applied after back-fitting microstate maps at global field power (GFP) peak locations only. Time points between GFP peaks are assigned the microstate label at the nearest GFP peak. As this method only permits microstate class switches half-way between GFP peaks, the method has a smoothing effect, compared to fitting every single time point. The name interpolation is justified due to the similarity to zero-order (‘sample and hold’) interpolation (Lyons, 2011). Both methods are widely used, but they have not been systematically compared.

As reviews and meta-analyses pool results across a large number of studies, the question arises to what extent heterogeneous preprocessing techniques might affect data variance and the final results. The question can be avoided as long as group differences within individual studies using a fixed pre-processing strategy are analyzed, i.e. when relative changes such as increased duration of a given microstate class under a certain condition are reported. However, the preprocessing and smoothing question becomes central when absolute or even normative parameter ranges are contrasted.

Koenig et al (2002) reported normative values for microstate durations across the lifespan using data from almost 500 subjects. Since all data were processed with the same method, preprocessing is not an issue, although different smoothing parameters would likely have resulted in different absolute values. A recent systematic review and meta-analysis extended this approach and reported normative ranges for duration and further temporal parameters across 93 microstate studies (Zanesco, 2024). Reviewing the studies entering this meta-analysis, as well as other review articles, reveals widely differing pre-processing approaches. The meta-analysis by Zanesco (2024) contains studies that used parametric smoothing with a total window size up to 86 ms (da Cruz et al, 2020), several studies using 62 ms (Jia and Yu, 2019; Pan et al, 2020; Zhang et al, 2021), and one study reporting a window size of 56 ms (Damborská et al, 2019), (D’Croz-Baron et al, 2019). Numerous other studies entering the meta-analysis used GFP peak interpolation (Milz et al, 2015, 2017; Schumacher et al, 2019; Li et al, 2021; Lian et al, 2021; Krylova et al, 2021; Coquelet et al, 2022; Liu et al, 2022). A third smoothing approach, often associated with the MATLAB software package by Poulsen et al (2018), enforces a minimum microstate duration and has been used in (Musaeus et al, 2019a,b, 2020; González-Villar et al, 2020; Hao et al, 2022). Some studies even combine interpolation and parametric smoothing (Gao et al, 2018). Lastly, more than a few studies fail to provide the information necessary to fully reproduce their microstate back-fitting approach. Several studies reporting extraordinarily long microstate durations of many hundreds of milliseconds do not provide back-fitting or smoothing details, which would be helpful for their interpretation (Cai et al, 2018, 2019; Bodda and Diwakar, 2025).

Overall, smoothing methods are expected to have marked effects on multiple temporal microstate parameters, but our understanding of this relationship is very limited. We hypothesized that smoothing does not merely rescale microstate parameters but can change the qualitative class of the inferred process, for instance, whether the process is memoryless or has long-range memory. To address this knowledge gap, this study analyzes smoothing effects on both elementary (duration, occurrence, coverage) and advanced dynamical metrics (syntax, short- and long-range correlations, complexity). Our aim is to clarify how methodological differences affect microstate statistics and to contribute toward more standardized and interpretable microstate research practices.

## Methods

### Experimental data

We analyzed resting-state EEG recordings from the LEMON database (Babayan et al, 2019). Pre-processed EEG files from the dataset contain continuous 61-channel EEG recordings with alternating eyes-open and eyes-closed conditions downsampled to 250 Hz (Babayan et al, 2019). This study analyzes eyes-closed data only. We split recordings at the indicated epoch boundaries and only retained eyes-closed epochs with a minimum duration of 58 seconds. We aimed for a minimum length of one minute but discovered that a slight reduction of this parameter led to the inclusion of considerably more EEG epochs. In total, we obtained 1383 eyes-closed epochs from N=201 subjects and each subject contributed either 6 or 7 epochs. Two subjects from the database were excluded because their epoch lengths did not meet our criteria.

### Pre-processing and microstate clustering

EEG filtering, microstate clustering, back-fitting and smoothing were all computed with the MNE Pycrostates package (Gramfort, 2013; Férat et al, 2022a). Individual EEG epochs were pre-processed by interpolating missing channels to obtain the full set of 61 EEG channels (Babayan et al, 2019), band-pass filtered to 1-30 Hz (zero-phase FIR filter, length 825 samples or 3.3 seconds, −6 dB cutoff frequencies at 0.50 Hz and 33.75 Hz), and an average reference was applied.

The global field power (GFP) time series σ_*t*_ records the standard deviation of the voltage vector across all EEG channels at each moment. Let 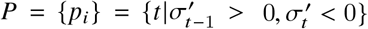 be the set of time points where the GFP attains local maxima (GFP peaks) and 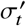 the first-order discrete derivative.

Voltage vectors were extracted at GFP peaks and submitted to modified K-means clustering with *K* = 5 clusters for each subject (Pascual-Marqui et al, 1995). Group-level microstate maps were computed from re-clustering the subject-level maps. The choice *K* = 5 was based on previous work on the same dataset where the range *K* = 1 … 12 was evaluated using the meta-criterion method (Férat et al, 2022b). We performed all computations for *K* = 4 and *K* = 6 as well (data not shown).

### Backfitting, sequence smoothing, and interpolation

Backfitting always used the winner-takes-all approach and the squared correlation coefficient between template maps and EEG data vectors as similarity measure. When no smoothing was applied (*b* = 0, or simply *b*0), the template maps were fitted to all available time points. We call this the raw microstate sequence.

#### Parametric smoothing

was introduced in Pascual-Marqui et al (1995) and uses a regularization method with two free parameters, the smoothing window half-size *b* ∈ N, and the strength (or Besag) factor λ ∈ R^+^. The algorithm starts with the raw microstate sequence and then proceeds iteratively, smoothing the microstate sequence until a convergence criterion is reached. Smoothing means that the algorithm contains a regularization term that penalizes microstate class switching within the temporal window [*t* − *b, t* + *b*] of size 2*b* + 1 samples. We varied *b* and used a constant strength parameter λ = 5. Further details are given in the supplementary material.

#### GFP peak interpolation

is another commonly used algorithm that starts with the sequence *L*_*t*_ defined at GFP peaks *t* ∈ *P* only. For the remaining time points, the microstate label at the nearest GFP peak is used. The full microstate sequence *X*_*t*_ is then defined as:

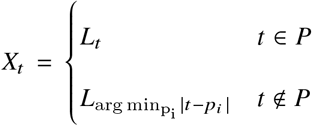

All pre-processing approaches yield a time series *X*_*t*_ over the categorical variable {0, 1, …, *K* − 1}. Using integer labels facilitates array indexing in subsequent analysis steps. When referring to the individual microstate classes, we label them *A*− *E*. We call these microstate sequences continuous sequences, since they have the same continuous time stamps as the underlying EEG. For each continuous sequence we obtain a jump sequence by removing consecutive duplicate labels. The continuous sequence snippet AACCCDDBAAA, for example, gives the jump sequence ACDBA (von Wegner et al, 2025). Jump sequences ignore the duration of a microstate, and only retain information about the transitions between the putatively underlying functional brain networks associated with each microstate. Microstate analysis computes duration and coverage from the continuous sequence and occurrence and transition matrices (first-order syntax) from the jump sequence. Other analyses can theoretically be computed on both sequences but are often more meaningful on only one of them. We will highlight which sequence type our analyses refer to.

### Elementary temporal metrics: duration, occurrence, coverage

The three elementary temporal metrics duration, occurrence and coverage were computed for each microstate class as described in the literature and implemented in all common microstate software packages.

### Markov tests

We conducted tests for the Markov property of order 0, 1 and 2 as described in earlier studies (von Wegner et al, 2017, 2025). The Markov test of order *m* tests whether the previous *m* + 1 time samples give a better prediction of the next future state than the previous *m* samples alone. If this is the case, the null hypothesis is rejected and provides evidence of time series memory extending over at least *m* + 1 samples. Thus, all three tests would be accepted for a Markov process of order 0. For a Markov process of order 1, the zero-order test would be rejected and the other two tests accepted. For a Markov process of order 2, the zero- and first-order tests would be rejected and the second-order test accepted. Higher-than-second-order processes would show all three tests being rejected. The following expressions are written in frequencies, or number of observations, denoted by the variable *f*. For *n* observations, we have *f*_*i*_ = *n* Pr(*X*_*t*_ = *i*), *f*_*ij*_ = *n* Pr(*X*_*t*−1_ = *i, X*_*t*_ = *j*), and so on.

Zero-order Markov test:

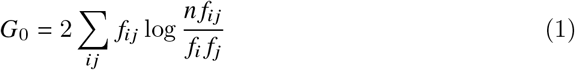

distributed as 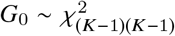.

First-order Markov test:

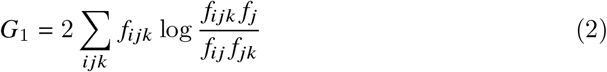

distributed as 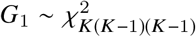.

Second-order Markov test:

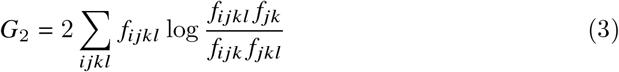

distributed as 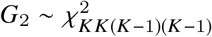.

### Transition matrix symmetry and entropy production

We investigated transition matrix symmetry and entropy production for continuous sequences as described before (Hermann et al, 2024; Manasova et al, 2025).

Transition matrix symmetry:

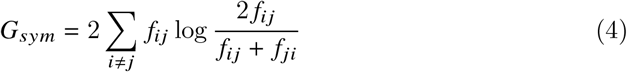

which is distributed as 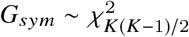 (von Wegner et al, 2017).

Entropy production quantifies time-irreversibility, or equivalently, the degree of detailed-balance violation of the microstate sequence and was calculated as Roldán and Parrondo (2010):

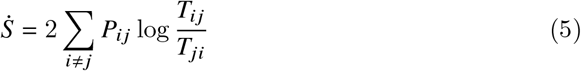

where *T*_*ij*_ = Pr(*X*_*t*+1_ = *j* | *X*_*t*_ = *i*) is the conditional transition probability and *P*_*ij*_ = *p*_*i*_ *T*_*ij*_ is the corresponding joint probability.

### Time scales

#### Relaxation time

Relaxation time describes how quickly the microstate sequence returns to its equilibrium distribution when disturbed. For a Markov process, this time constant is encoded in its transition matrix. Non-Markovian processes can deviate considerably from this behaviour but we have found close relationships between the relaxation time, microstate durations, and entropy rates in the past (Wiemers et al, 2023). The relaxation time of each microstate sequence is computed from the eigenvalues of its transition matrix *T*_*ij*_ defined above (von Wegner and Laufs, 2018)

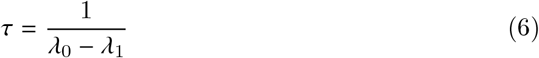

where λ_0_ = 1 is the largest, and λ_1_ the second-largest eigenvalue.

#### Autoinformation function

The autoinformation function (AIF) measures two-point correlations between time points separated by a fixed time lag and can be regarded as a generalized autocorrelation function. We arbitrarily defined intermediate time scales as lags up to 400 ms, covering the main EEG frequency bands. AIF peaks indicate periodic features in the underlying microstate sequence (von Wegner et al, 2017). The autoinformation coefficient *I*(*k*) for time lag *k* (samples) is the time-lagged mutual information between *X*_*t*_ and *X*_*t*+*k*_ :

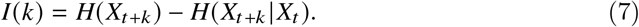

where *H*(·) denotes the Shannon entropy of a random variable.

#### Long-range correlations

We investigated the long-range behaviour by calculating a Hurst exponent *H* for each microstate sequence using a random walk representation (Van de Ville et al, 2010; von Wegner et al, 2016; Jia et al, 2021). We calculated *H* with detrended fluctuation analysis (DFA) as described before (von Wegner et al, 2016; Jia et al, 2021), averaging over the ten possible (2, 3) partitions for *K* = 5, i.e. ((*A, B*), (*C, D, E*)), ((*A, C*), (*B, D, E*)), …. Hurst exponents were fitted to the detrended fluctuations across 50 different window sizes log-uniformly spaced between minimum and maximum scales of 150 samples (600 ms) and 3000 samples (12 seconds), respectively.

### Complexity and higher-order syntax

For continuous microstate sequences, we computed their entropy rate and excess entropy as metrics of type I and type II complexity, respectively (Sleigh and Hight, 2021). The implementation was identical to (von Wegner et al, 2023).

The structure of this study is summarized in Figure 1. While elementary temporal metrics and Markov tests are computed on both the continuous and jump sequences, the extended autocorrelation structure is studied on continuous sequences only. The time scale distinction into tens, hundreds and thousands of milliseconds for relaxation time, AIF, and Hurst exponents, respectively, is approximate. AIF time lags start at zero and thus contain the same short time lags that Markov tests address, and there is significant overlap between AIF and Hurst exponent time scales.

**Fig. 1.**
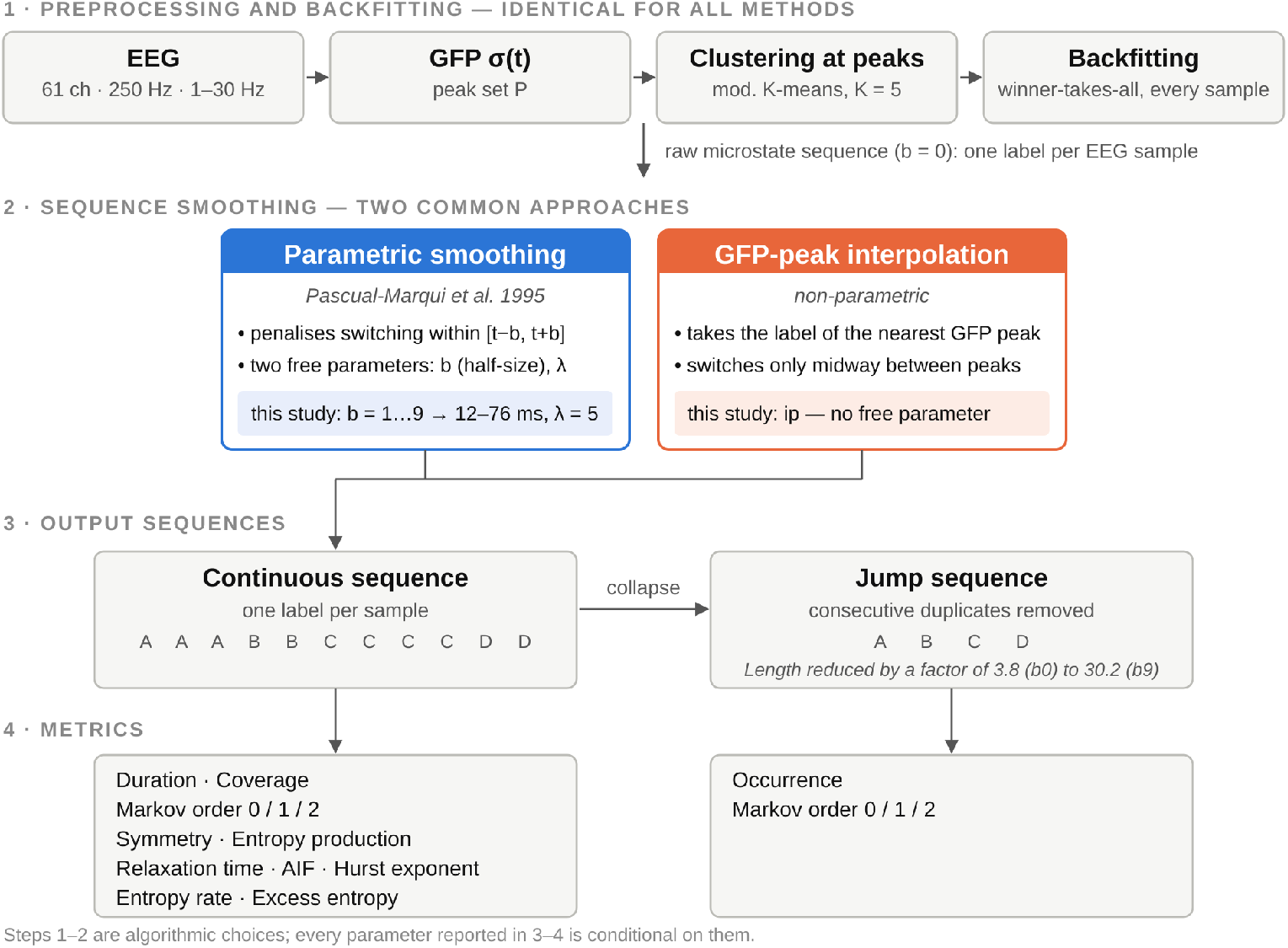
Overview: We quantify the effects of two different smoothing algorithms on a range of metrics characterizing microstate sequences. Pre-processing and back-fitting follow established microstate processing standards (1), followed by either windowed parametric smoothing or GFP peak interpolation (2). Parametric smoothing is evaluated for window half-size values of *b* = 1, 3, 5, 7, 9, corresponding to total window sizes of 12, 28, 44, 60, 76 ms. Some metrics are evaluated on continuous microstate sequences containing state duration information, others on jump sequences that only record state transitions (3,4).

### Statistics

All metrics were analyzed across the factor ‘smoothing method’ which had seven levels, representing parametric smoothing with strength λ = 5 and window half-size *b* = 0 (no smoothing), *b* = 1, 3, 5, 7, 9 samples, and ip for GFP peak interpolation. The results were considered a set of repeated measures since the same microstate sequences were processed by different smoothing procedures.

All parameters except symmetry tests were compared with the non-parametric Friedman test 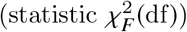 and the effect size was recorded as Kendall’s *W*. When significant (*p* < 0.05), we conducted post-hoc pairwise Nemenyi tests. All significance levels were set to α = 0.05.

For Markov and symmetry tests, we retained the test results as p-values, which were pooled across all epochs for each subject via the harmonic mean and corrected for multiple comparisons (Benjamini-Hochberg FDR). For the symmetry test, the fraction of rejected null hypotheses is a binary per-subject outcome under repeated measures and was tested with Cochran’s Q test at α = 0.05, followed by post-hoc pairwise McNemar tests (FDR-corrected).

## Results

### Effects on sequence length

Smoothing has minimal effects on the length of continuous sequences. The only mechanism that changes the length of a continuous sequence is the removal of the first and the last microstate in a sequence, due to their unknown length. Since smoothing increases the duration of microstates, more samples are removed when the smoothing window is wider. The first column of Table 1 shows that continuous sequences are usually shortened by less than 100 samples, relative to an average total length of more than 15,000 samples. Jump sequences, however, are shortened significantly (columns 3 and 4 in Table 1). Without smoothing, jump sequences have, on average, 3.76 times fewer samples than the corresponding continuous sequence. The mean shortening factor increases to 30.23 for a 76 ms smoothing window (b9). The average shortening factor for GFP peak interpolation (ip) was 13.83, comparable to smoothing windows of 28-44 ms (b3-b5).

**Table 1.**
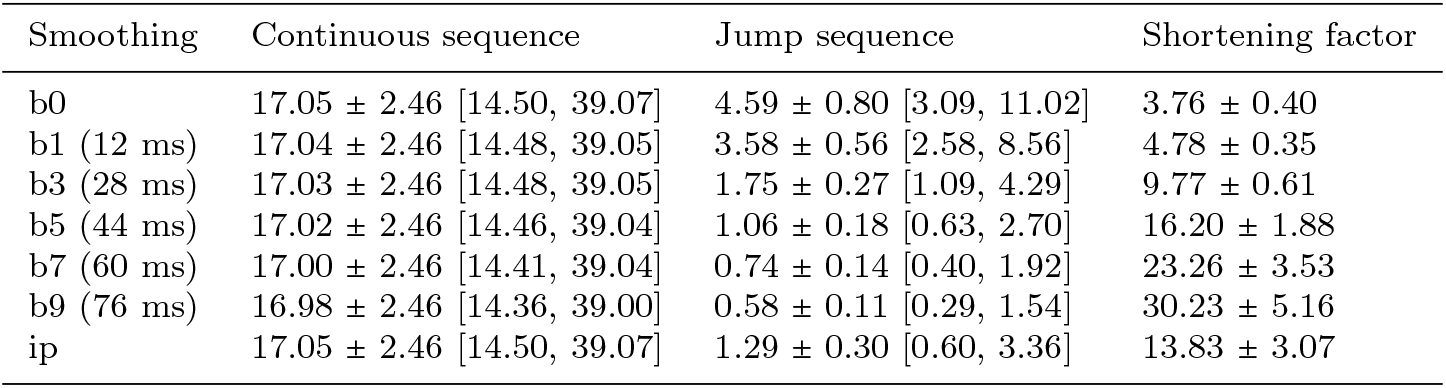
Microstate sequence length as a function of sequence smoothing. Sequence lengths are given in kilosamples (1000 samples), as mean ± standard deviation [min, max]. One minute of EEG corresponds to 15000 samples (250 Hz). *b*0: no smoothing, *b*1 ™ *b*9: parametric smoothing, ip: interpolation between GFP peaks.

| Smoothing | Continuous sequence | Jump sequence | Shortening factor |
| --- | --- | --- | --- |
| b0 | 17.05 ± 2.46 [14.50, 39.07] | 4.59 ± 0.80 [3.09, 11.02] | 3.76 ± 0.40 |
| b1 (12 ms) | 17.04 ± 2.46 [14.48, 39.05] | 3.58 ± 0.56 [2.58, 8.56] | 4.78 ± 0.35 |
| b3 (28 ms) | 17.03 ± 2.46 [14.48, 39.05] | 1.75 ± 0.27 [1.09, 4.29] | 9.77 ± 0.61 |
| b5 (44 ms) | 17.02 ± 2.46 [14.46, 39.04] | 1.06 ± 0.18 [0.63, 2.70] | 16.20 ± 1.88 |
| b7 (60 ms) | 17.00 ± 2.46 [14.41, 39.04] | 0.74 ± 0.14 [0.40, 1.92] | 23.26 ± 3.53 |
| b9 (76 ms) | 16.98 ± 2.46 [14.36, 39.00] | 0.58 ± 0.11 [0.29, 1.54] | 30.23 ± 5.16 |
| ip | 17.05 ± 2.46 [14.50, 39.07] | 1.29 ± 0.30 [0.60, 3.36] | 13.83 ± 3.07 |

### Elementary temporal metrics

Smoothing effects on duration, occurrence and coverage are shown in Figure 2, and the numerical results are summarized in Table 2. Friedman tests across the factor’smoothing method’ were significant for all three parameters. Window half-size *b* increased microstate durations (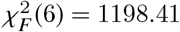, *p* < 0.001) and had a large effect size (*W* = 0.994). All pairwise post-hoc comparisons were significant. GFP peak interpolated sequences had durations between those for smoothing window sizes 28 ms (b3) and 44 ms (b5).

**Table 2.** Duration, occurrence, and coverage for microstate classes A-E across smoothing methods. All values are given as mean (standard deviation).

| smoothing | ms-class | Duration (ms) | Occurrence (Hz) | Coverage |
| --- | --- | --- | --- | --- |
| b0 | A | 13.61 (1.67) | 13.34 (2.27) | 0.18 (0.04) |
|  | B | 14.38 (1.77) | 13.88 (2.06) | 0.20 (0.03) |
|  | C | 16.63 (3.41) | 13.91 (1.63) | 0.23 (0.06) |
|  | D | 13.85 (2.07) | 12.72 (1.93) | 0.18 (0.04) |
|  | E | 15.65 (2.33) | 13.54 (1.91) | 0.21 (0.04) |
| b1 (12 ms) | A | 17.77 (1.63) | 10.19 (1.67) | 0.18 (0.04) |
|  | B | 18.50 (1.69) | 10.76 (1.43) | 0.20 (0.04) |
|  | C | 20.42 (3.38) | 11.13 (1.54) | 0.23 (0.06) |
|  | D | 17.81 (2.08) | 9.78 (1.54) | 0.18 (0.04) |
|  | E | 19.64 (2.28) | 10.75 (1.40) | 0.21 (0.04) |
| b3 (28 ms) | A | 36.40 (2.74) | 4.74 (0.98) | 0.17 (0.05) |
|  | B | 37.65 (3.11) | 5.18 (0.79) | 0.20 (0.04) |
|  | C | 41.96 (7.94) | 5.72 (0.97) | 0.25 (0.09) |
|  | D | 35.30 (2.81) | 4.57 (1.04) | 0.16 (0.05) |
|  | E | 38.98 (4.20) | 5.50 (0.83) | 0.22 (0.06) |
| b5 (44 ms) | A | 58.82 (5.43) | 2.87 (0.72) | 0.17 (0.05) |
|  | B | 61.48 (6.78) | 3.14 (0.60) | 0.20 (0.05) |
|  | C | 70.88 (19.25) | 3.49 (0.59) | 0.25 (0.10) |
|  | D | 57.34 (5.70) | 2.77 (0.74) | 0.16 (0.05) |
|  | E | 64.18 (9.12) | 3.36 (0.61) | 0.22 (0.06) |
| b7 (60 ms) | A | 82.92 (9.06) | 2.01 (0.56) | 0.17 (0.06) |
|  | B | 87.24 (11.61) | 2.21 (0.48) | 0.19 (0.06) |
|  | C | 102.98 (33.30) | 2.45 (0.44) | 0.26 (0.11) |
|  | D | 80.91 (9.70) | 1.95 (0.58) | 0.16 (0.06) |
|  | E | 91.46 (15.46) | 2.37 (0.49) | 0.22 (0.07) |
| b9 (76 ms) | A | 106.35 (12.52) | 1.55 (0.46) | 0.17 (0.06) |
|  | B | 112.37 (16.81) | 1.71 (0.40) | 0.19 (0.06) |
|  | C | 134.61 (47.70) | 1.90 (0.36) | 0.26 (0.11) |
|  | D | 103.49 (13.40) | 1.51 (0.47) | 0.16 (0.06) |
|  | E | 118.35 (22.12) | 1.83 (0.41) | 0.22 (0.07) |
| ip | A | 47.98 (6.65) | 3.50 (1.09) | 0.16 (0.05) |
|  | B | 51.34 (8.51) | 3.85 (0.97) | 0.19 (0.05) |
|  | C | 65.23 (23.12) | 4.48 (0.67) | 0.29 (0.09) |
|  | D | 45.59 (6.65) | 3.00 (0.99) | 0.13 (0.04) |
|  | E | 54.43 (12.02) | 4.08 (0.95) | 0.22 (0.06) |

**Fig. 2.**
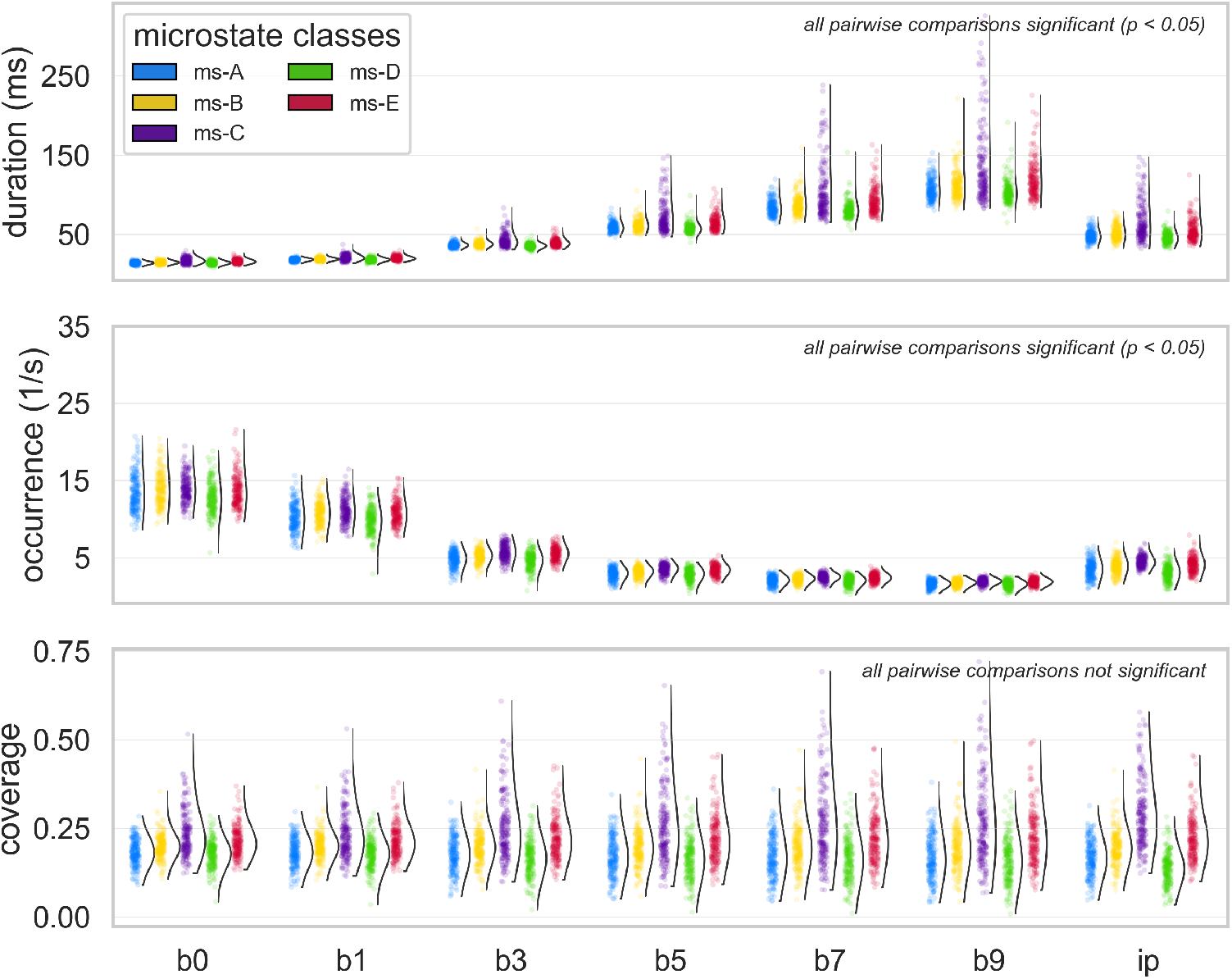
Effects of microstate sequence smoothing on duration (top), occurrence (middle), and coverage (bottom) for *K* = 5 classes (A-E). Raincloud plots show the individual data points for N=201 subjects in the left half, and the corresponding density plot in the right half. For each subject, each microstate parameter has been averaged across the available epochs for that subject. Friedman tests were positive (*p <* 0.05) for all three parameters. For duration and occurrence, all pairwise differences between smoothing methods were significant, none was significant for coverage.

Microstate occurrence decreased with wider smoothing windows (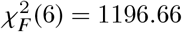, *p* < 0.001), again with large effect size (*W* = 0.992). All pairwise post-hoc comparisons were significant. GFP peak interpolated sequences had occurrence rates between those for smoothing window sizes 28 ms (b3) and 44 ms (b5).

The omnibus test for microstate coverage across smoothing methods was significant with very small effect size (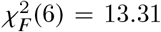, *p* = 0.038, *W* = 0.011) but none of the pairwise comparisons was significant.

### First-order syntax and Markov properties

As smoothing affects the temporal order of microstates in a sequence, we tested whether low-order Markov properties would be affected by the different smoothing approaches. Figure 3 shows the results of low-order Markovianity tests for continuous microstate sequences on the left, and for jump sequences on the right. The abscissa indicates the smoothing method and the ordinate indicates the fraction of accepted Markovianity tests, *Pr* (*p*_*FDR*_ ≥ 0.05), where p-values were FDR-corrected.

**Fig. 3.**
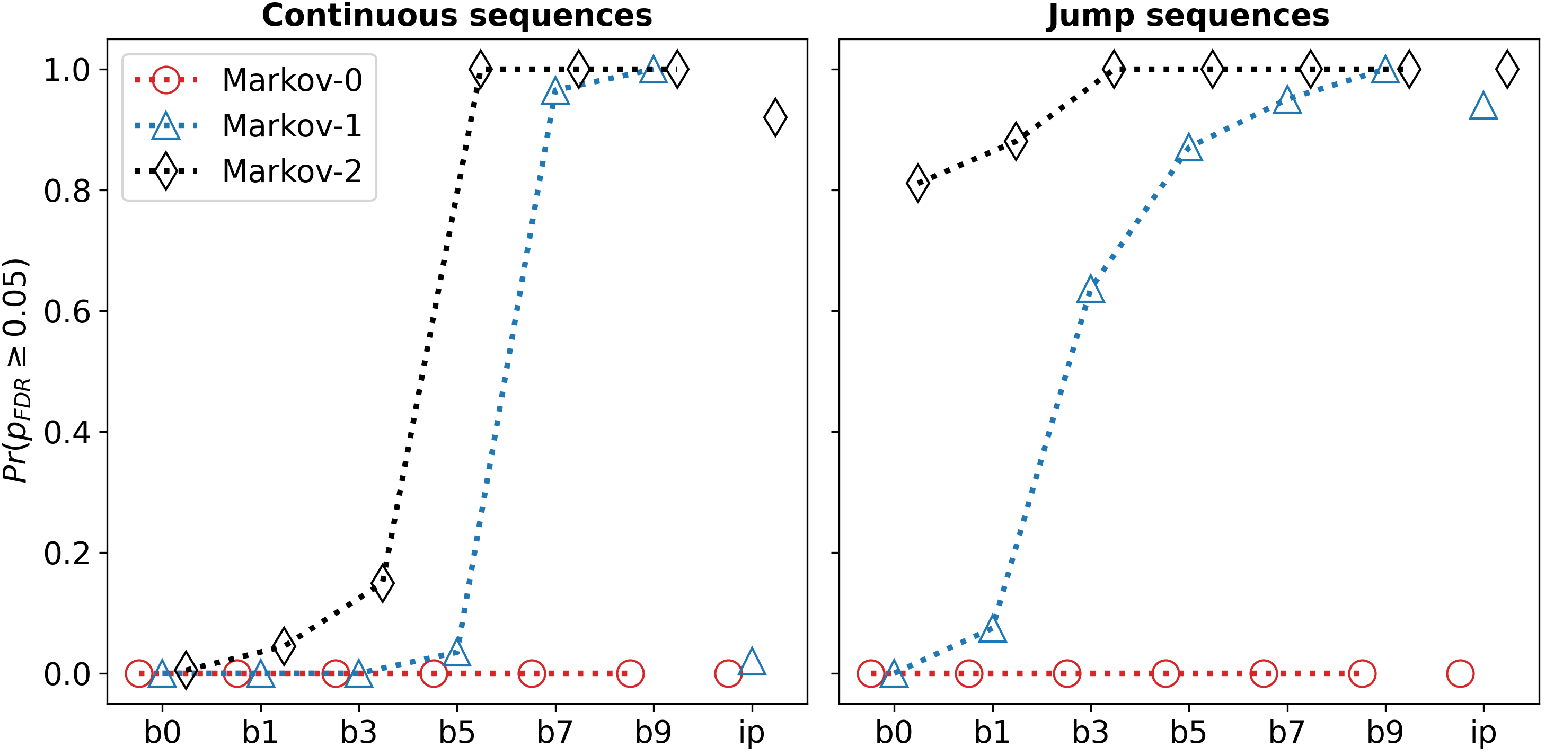
Markov properties of continuous (left) and jump (right) microstate sequences across smoothing methods. The ordinate shows the fraction of accepted Markov tests. Smoothing affects the apparent Markov order of microstate sequences by progressively eliminating memory effects at short time scales.

For continuous sequences, all zero-order tests are rejected. Jump sequences cannot have order zero by definition. The first-order property is fully accepted up to *b* = 3 for continuous sequences and for *b* = 0 jump sequences. The second-order property is fully rejected at *b* = 5 for continuous sequences and at *b* = 3 for jump sequences. Overall, stronger smoothing (larger value of *b*) suppresses higher-order Markov properties, eventually giving all sequences a first-order appearance. GFP peak interpolation had effects comparable to a smoothing window half-size of *b* = 3 − 5 (28-44 ms).

### Transition matrix symmetry and entropy production

The fraction of subjects with a significant departure from conditional transition symmetry (*p* < 0.05) dropped sharply with smoothing, from 89% (*b*0), 76% (*b*1) and 11% (*b*3) to ≤2% for *b*5 and all wider windows, as shown in the upper half of Figure 4. Effects were highly significant (Q(6) = 892.20, *p* < 0.001). Post-hoc pairwise McNemar tests (FDR-corrected) showed that *b*0, *b*1, and *b*3 differed significantly from every other condition and from each other (all *p* < 0.001), whereas *b*5, *b*7, *b*9, and *ip* were mutually indistinguishable (all corrected *p* ≥ 0.62).

**Fig. 4.**
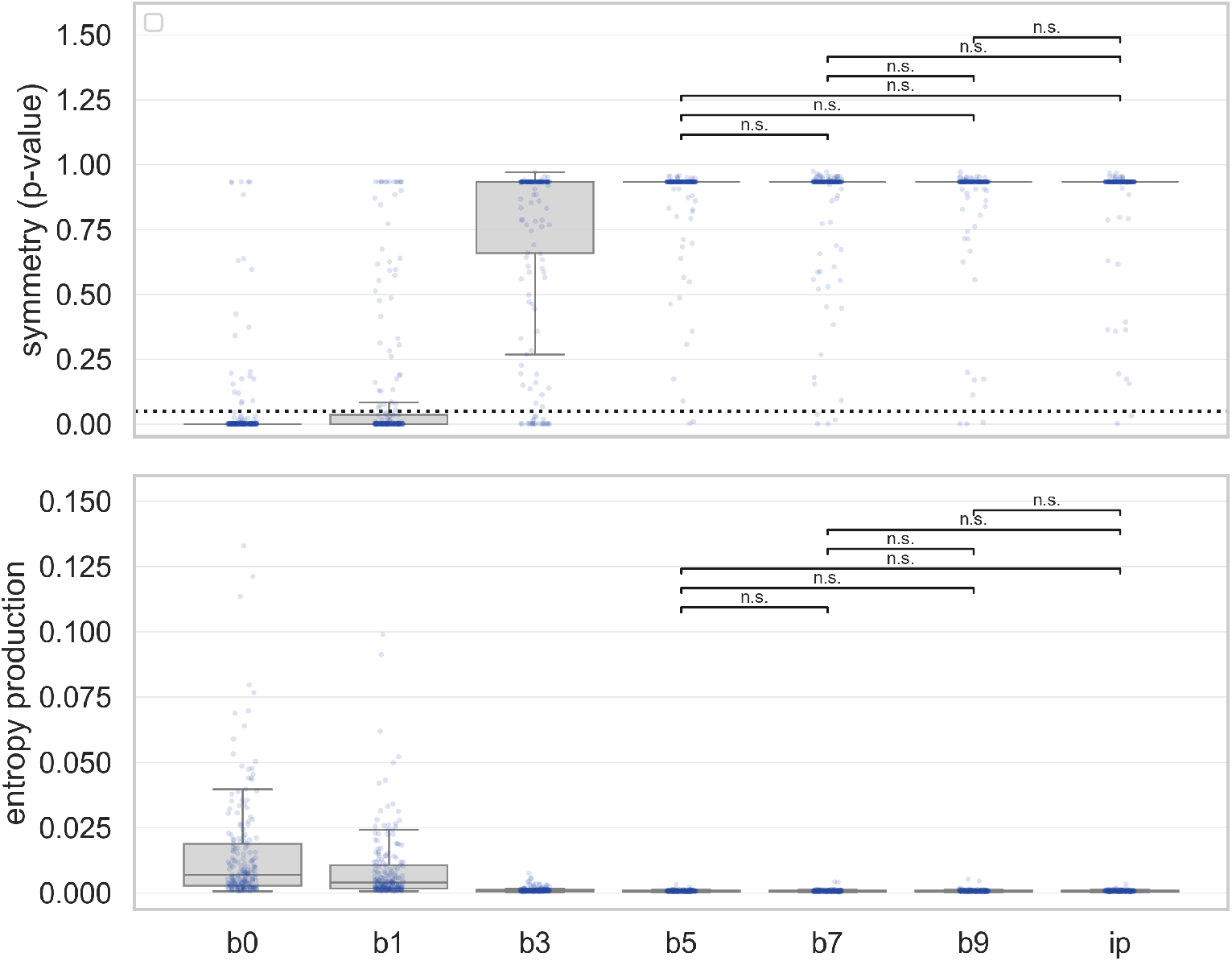
Transition matrix properties (continuous sequences). Top: Results of the conditional transition probability matrix symmetry test. The y-axis indicates p-values of test results across all subjects (harmonic mean across epochs per subject). The dashed line is set at *p* = 0.05 and *p <* 0.05 indicates transition matrix asymmetry. Bottom: Entropy production statistics. Values larger than zero indicate positive entropy production. In both panels, only non-significant (n.s., *p >* 0.05) pairwise differences are shown for better visibility.

Entropy production (or detailed balance), shown in the lower half of Figure 4, decreased monotonically with window size. A Friedman test confirmed a significant overall effect (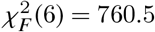, *p* < 0.001) with a large effect size (Kendall’s *W* = 0.631). Post-hoc Nemenyi tests showed the same pattern as for symmetry: *b*0, *b*1 and *b*3 differed significantly from all other conditions and from each other, while *b*5, *b*7, *b*9, and *ip* formed a single non-significant cluster (all pairwise *p* ≥ 0.90).

### Intermediate and long time scales

In this section, we report the effects of smoothing on short, intermediate and long time scales. The results are shown in Figure 5.

**Fig. 5.**
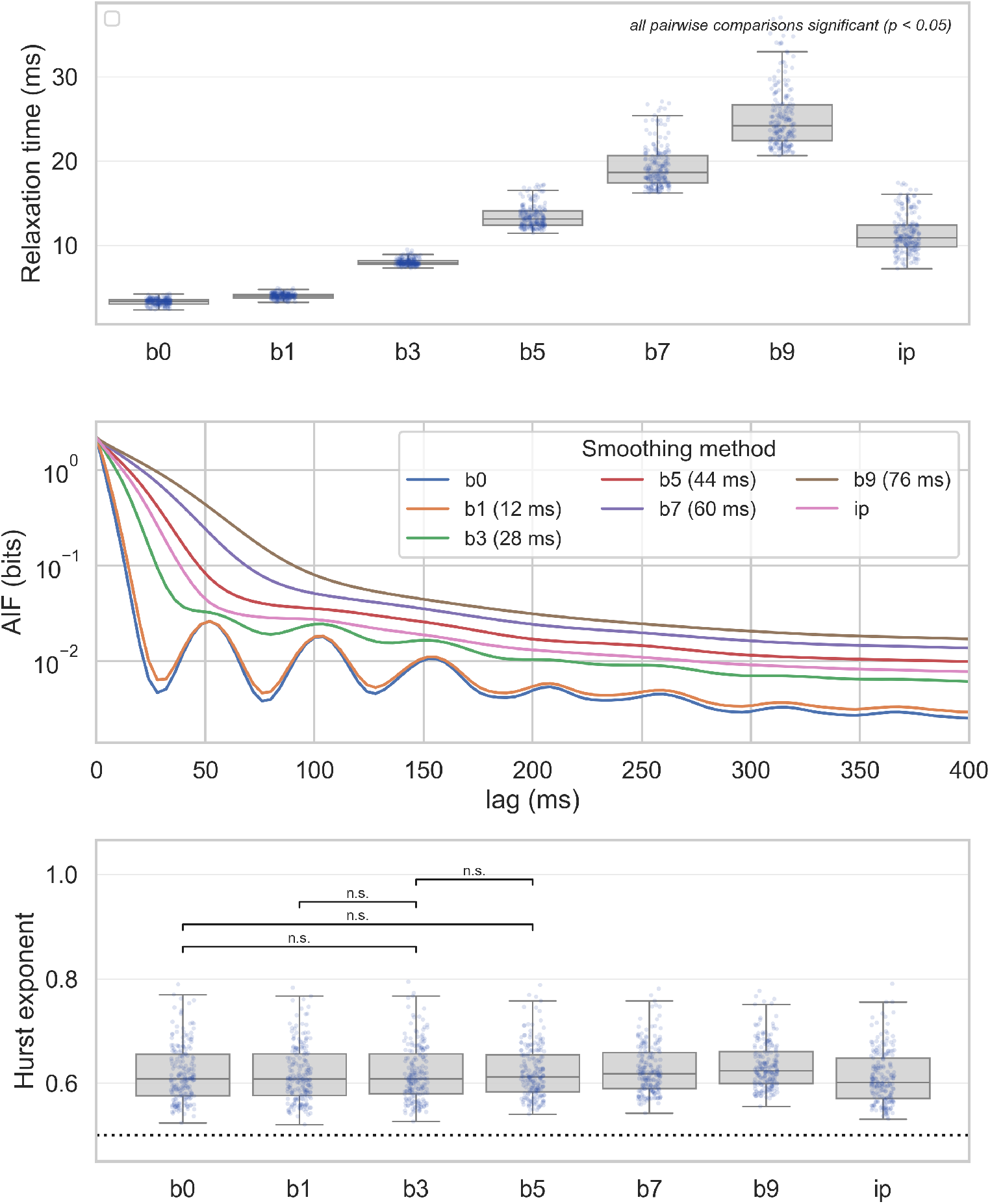
Effects of microstate sequence smoothing on different time scales inherent to continuous microstate sequences. Top: The transition matrix relaxation time increases with smoothing parameter *b*. All pairwise comparisons were statistically significant (*p* < 0.001). Middle: Autoinformation functions (AIF). Mutual information for all time lags increases with smoothing intensity. Periodicity is observed up to a smoothing window half-size of *b* = 3. Bottom: Hurst exponents obtained from detrended fluctuation analysis (DFA). The absence of long-range correlations is indicated by *H* = 0.5 (dashed line). For all smoothing methods, persistence (*H* > 0.5) was confirmed. Non-significant (n.s., *p >* 0.05) pairwise differences are indicated.

#### Relaxation time

The effects of different smoothing methods on the relaxation time are shown in the top panel of Figure 5. The main effect of smoothing was significant (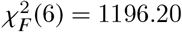, *p* < 0.001) with large effect size (*W* = 0.992). Post-hoc testing showed that all 21 pairwise comparisons between smoothing methods were significant (*p* < 0.05).

#### Intermediate time scales

We assessed the time scale range from tens to hundreds of milliseconds by the AIF. The results are shown in the middle panel of Figure 5 and several effects can be observed. First, the AIF curves of microstate sequences that have undergone more intense smoothing lie above the AIF curves of less smoothed sequences. Periodicities are observed for *b*0 (no smoothing), *b*1 (12 ms) and *b*3 (28 ms). Beyond that, and including the interpolation method, sequence periodicity cannot be observed. Also, smoothing affects the lower end of the AIF curve (up to 100 ms) more than the far end where time lags of up to 400 ms are represented. The AIF of GFP peak interpolated sequences (pink curve) lies between the *b*3 and *b*5 curves.

#### Long-range correlations

We quantified long-range correlations (time scales 600 ms - 12 s) by the Hurst exponent, the results are shown in the bottom panel of Figure 5. Smoothing had significant effects (χ^2^ (6) = 561.03, *p* < 0.001) with moderate effect size (*W* = 0.465). Across smoothing methods, the group mean values ranged from *H* = 0.61 to *H* = 0.63. Post-hoc pairwise testing showed non-significant results between *b*0 − *b*3, *b*0 − *b*5, *b*1 − *b*3, and *b*3 − *b*5; all other comparisons were statistically significant.

For each smoothing method, Hurst exponents were significantly larger than *H* = 0.5 (one-sided Wilcoxon test, *p* < 0.001 for all smoothing methods), indicating persistence.

### Sequence complexity and higher-order syntax

Smoothing effects on information-theoretic parameters are shown in Figure 6. Type I complexity (entropy rate) decreased with stronger smoothing (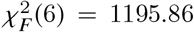, *p* < 0.001) with large effect size (*W* = 0.992). All 21 pairwise post-hoc comparisons were statistically significant. Type II complexity (excess entropy) increased with stronger smoothing (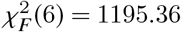, *p* < 0.001) with large effect size (*W* = 0.991). All 21 pairwise post-hoc comparisons were statistically significant. Both entropy rate and excess entropy values for GFP peak interpolated microstate sequences were similar to those calculated with smoothing windows of size 28 ms (b3) and 44 ms (b5).

**Fig. 6.**
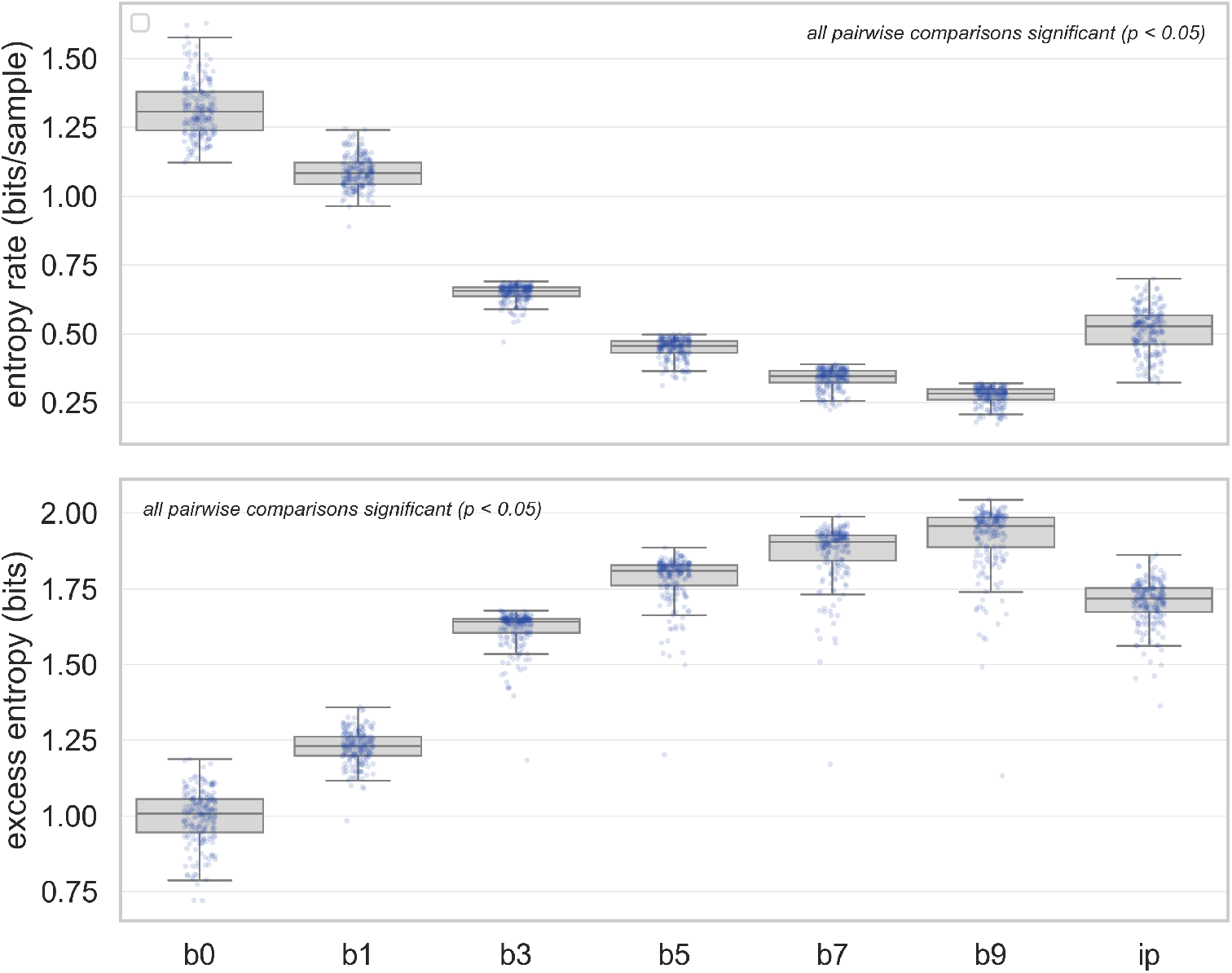
Effects of microstate sequence smoothing on type I and type II complexity of continuous microstate sequences. Top: Entropy rate decreases with more intense smoothing. All pairwise comparisons were statistically significant. Bottom: Excess entropy increases with smoothing. All pairwise comparisons were statistically significant.

## Discussion

This study highlights the impact of temporal smoothing on EEG microstate sequences, and how algorithmic and parameter choices result in different perspectives on resting-state brain activity. The main findings are that sequence smoothing

1. leads to a continuous change in microstate duration and occurrence, but not coverage;
2. at short time scales, gives microstate sequences the structure of a memory-less (Markovian), symmetric, near-equilibrium process with increasingly slower relaxation time;
3. removes periodic features from microstate sequences;
4. has minor effects on long-range correlations and the Hurst phenomenon;
5. decreases type I (entropy rate) and increases type II (excess entropy) complexity metrics, and
6. using a smoothing window (total) size of 30-40 ms has effects similar to GFP peak interpolation.

### Elementary temporal metrics

As expected, sequence smoothing had marked effects on the parameters duration and occurrence. As these parameters are direct consequences of where the microstate boundaries are located, smoothing increases duration and reduces occurrence rates. Effects on coverage were minor. Assuming constant coverage, the previous observation is then easily explained by the relationship: occurrence ∝ duration^-1^.

Using a smoothing parameter range taken from the literature, we observed a wide range of microstate durations. Microstate durations are among the most frequently reported parameters and are interpreted as a measure of temporal stability of the functional neuronal assembly associated with the given microstate map (Michel and Koenig, 2018). One argument for the functional relevance of microstates is the overlap of their duration with the time scales of perception and cognitive processing (Michel and Bréchet, 2026). While this suggests that these time scales are rediscovered during microstate analysis, our results show that the choice of preprocessing parameters might have a much stronger impact than assumed. Some authors even state that “parameter settings were determined through multiple repetitive tests to ensure that the average duration of microstates was in the range of 80-150 ms” (Shi et al, 2026), making microstate durations a tunable property of microstate analysis rather than an objective outcome. Empirically, microstate durations increase monotonically with smoothing window half-size *b*, without forming a plateau that might suggest a fixed underlying time scale. Mean durations in our study varied more than five-fold, covering a range of values from less than 20 ms to more than 100 ms across the tested parameter range. Microstate analysis does not come with an internal validation procedure that could identify the true duration, if it exists. Strictly speaking, microstate durations should only be compared when at least similar preprocessing techniques have been applied. Vice versa, similar microstate durations obtained under significantly different smoothing parameters should rather be considered non-identical. Given the continuous increase in duration, and decrease in occurrence, our analyses do not allow a conclusion regarding the true value of these parameters. Further studies combining traditional microstate analysis with invasive EEG might help gaining further insights (Mikutta et al, 2024).

A side effect of smoothing is a significant shortening of jump sequences obtained from the initial continuous sequences by collapsing duplicates into single labels. We observed shortening factors of up to 30 for the largest smoothing windows, resulting in microstate sequences as short as a few hundred samples per one minute of EEG data. This has an impact on the statistical power of standard syntax tests using χ^2^ tests (Lehmann et al, 2005) and even more so on data-hungry methods assessing higher-order properties and complexity metrics where short sequence lengths cause false negatives and biased estimators (von Wegner et al, 2023, 2025, 2026).

### Short time scales and syntax properties

Time scales similar to the smoothing window size are directly altered by smoothing. While this has the desired effect of removing short blocks of labels, that are neuro-physiologically incompatible with large-scale network reconfigurations, it also imposes certain syntax properties on microstate sequences. Markovianity tests (Figure 3) show that smoothing windows between 28 ms and 44 ms cause a nearly complete transition to a first-order Markov law and that the effect occurs earlier in jump sequences. Additional simulations not contained in the manuscript showed that the jump sequence Markov tests were not accepted due to the shorter sequence length which should only occur when the sequences become as short as a few hundred samples. As discussed below, the apparent first-order Markov structure detected at short time lags does not remove long-memory, persistent features documented by the Hurst phenomenon, resulting in an overall inconsistent picture of memoryless Markovian dynamics when looking at short time scales and non-Markovian long-memory effects when inspecting large time scales.

Similarly, as shown in Figure 4, conditional and joint transition probabilities appear increasingly symmetric. The latter observation represents loss of time-irreversibility and suggests a state closer to thermodynamic equilibrium (Lynn et al, 2021). Yet, current research involving microstate analysis (Manasova et al, 2025), electrocorticography in animals (de la Fuente et al, 2023), and fMRI (Lynn et al, 2021), suggests that time-irreversibility and non-equilibrium brain dynamics are hallmarks of intact consciousness and correlate positively with cognitive load. This would suggest that the fingerprints of non-equilibrium dynamics found for short smoothing windows (*b* = 1 or 12 ms) in this study might reflect a meaningful neurophysiological signal. It is not caused by single intermittent microstate labels as these are removed even with short smoothing windows. These findings speak in favour of shorter smoothing windows.

### Intermediate time scales and periodic microstate features

Smoothing also affected time scales beyond the first few samples. Relaxation time is a global measure (not per microstate) of how fast a microstate sequence returns to its equilibrium distribution and covaries closely with individual microstate mean durations and entropy rate (Wiemers et al, 2023). This parameter also increased continuously with stronger smoothing, resulting in an image of globally slower brain dynamics.

Our analysis confirms that microstate sequences show periodic features not explained by single noisy states near GFP troughs as these are eliminated by short smoothing windows. Periodic features persisted up to a window length of 28 ms (b=3) in Figure 5 but quickly disappear beyond time scales approaching the interval between GFP peaks. This might explain why some authors fail to identify periodic features (Al Zoubi et al, 2019). Based on our findings, microstate durations around 60 ms as reported by the authors suggest that either GFP peak interpolation or smoothing with a window size of approximately 40 ms might have been used. In a previous article, we found that spectral effects of low-dose propofol could be observed in non-smoothed but not in interpolated sequences (Hermann et al, 2024). The results of this study emphasize that microstates between GFP peaks are not random switches caused by the absence of a recognizable electric field configuration, or simply ‘noise’, as these would have been eliminated by smoothing windows spanning seven samples (b3). The b3 filter preserves periodic microstate features but spans 28 ms, more than half the common GFP peak-to-peak interval during eyes-closed resting state. While it is true that the variance of the field goes through local troughs, many of these minima have higher GFP values than other GFP peaks, and winner-takes-all fitting still reveals highly structured transitions between GFP peaks. As these periodic features potentially encode meaningful information, we suggest smoothing windows well below the GFP peak-to-peak interval.

### Persistence and long-range correlations

The effect of smoothing on Hurst exponents was mixed. While there was a significant overall effect, persistence (*H* > 0.5) was reliably detected for all settings. The effect size was moderate and numerous pairwise differences were not statistically significant, especially in the range *b*0 − *b*5. The overall effect was somewhat surprising since the largest smoothing windows were 76 ms and the DFA fitting range started at 600 ms in our study. This might suggest a minor role of shorter time scales on individual *H* estimates. Overall, the Hurst exponent can be regarded as a stable parameter, facilitating its comparison across studies. The origin of the Hurst phenomenon in microstate sequences, however, is still being investigated. Slow exponential time scales and non-stationarities such as vigilance fluctuations are likely to contribute.

### Complexity

The complexity metrics considered, entropy rate for type I and excess entropy for type II complexity, were strongly affected by smoothing, decreasing type I and increasing type II complexity. This finding is explained by the fact that the actual microstate word size used overlapped with the size of smoothing windows. As these metrics require estimates of multivariate distributions, the meaningful word size is usually limited to less than 10 labels. Even if this was not the case, we know that these metrics are influenced by genuine EEG slowing as it occurs during non-REM sleep (Wiemers et al, 2023; von Wegner et al, 2023) and that this effect is mediated by first-order, not higher-order, dependencies (von Wegner et al, 2023). Nevertheless, these metrics are capable of capturing higher-order interactions and differentiating clinical populations (von Wegner et al, 2026). The global effect of smoothing can be described as reduced randomness (type I) and increased information transfer from past to future states, which is due to the pressure to homogenize states inside the smoothing window.

### Conclusions and recommendations

Considering neurophysiological constraints of large-scale network activity, removing microstates lasting less than 20 ms is reasonable. On the other hand, our results imply that strong smoothing might attenuate or even remove meaningful biomarkers of normal brain activity. Smoothing can induce inconsistent results across time scales, namely memoryless patterns at short time scales vs. long-range memory features at long time scales, remove signs of non-equilibrium dynamics and periodic features. A window length of no more than 30 ms seems adequate for resting-state wakefulness in healthy subjects. Care needs to be taken when setting these parameters as software parameters sometimes ask for the *window half-size*. GFP peak interpolation seems to be slightly beyond this safety margin and does remove some of the dynamic features listed above. Any smoothing procedure should be documented precisely. Comparative analyses including meta-analyses should take pre-processing strategies into account.

## Acknowledgment

We thank Victor Férat for discussions of smoothing algorithms in Pycrostates.

## Declarations

This work was funded by the German Research Foundation (DFG), project number 440536202.

## Supplemental Data

### Parametric smoothing

We followed the algorithm given in Pascual-Marqui et al (1995). The procedure starts with the raw microstate sequence and then proceeds iteratively until a convergence criterion is reached. The algorithm contains a regularization term that penalizes microstate class switching within a temporal window [*t* −*b, t* + *b*]. During each iteration, the following function is minimized:

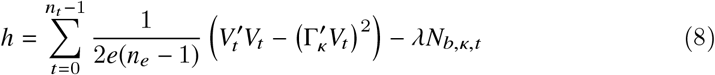

Where

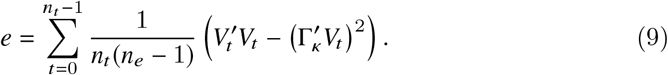

is an estimate of the residual variance after the initial fit and is computed once before the iteration starts.

Using the notation from the original article Pascual-Marqui et al (1995):

- *V*_*t*_ is the voltage (column) vector, an *n*_*e*_-channel EEG sample, taken at time *t* from a total of *n*_*t*_ samples. 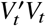 is the scalar product of *V*_*t*_ with itself, i.e. the squared norm ∥*V*_*t*_ ∥ ^2^.
- κ = *X*_*t*_ is the current microstate label at time *t*, which can change at each iteration.
- 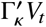 is the scalar product of the best-matching microstate map Γ_κ_ from the previous iteration and the current voltage vector *V*_*t*_ and indicates their degree of similarity. We have 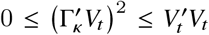 and minimization of this term favours a high correlation between the microstate map Γ_κ_ and the current data vector.
- *N*_*b*,κ,*t*_ is the number of microstate sequence labels equal to κ within the discrete neighbourhood [*t* − *b, t* + *b*]. Due to the negative sign, optimization tries to maximize this term, thus favouring contiguous blocks of microstate label κ within the current smoothing window. Note that Pascual-Marqui et al (1995) exclude the label at *t* while the frequently used software packages described in Nagabhushan Kalburgi et al (2023) and Férat et al (2022a) use a convolution operator that includes *t*. As many results reported in the literature would have been produced with convolution, we also opted for this method.
- λ is a free parameter weighing the influence of *N*_*b*,κ,*t*_ and is often called the strength parameter or the Besag factor (Besag, 1986). Larger values lead to stronger smoothing without changing the window size.

#### The strength parameter λ

The second free parameter, denoted λ in the original article (Pascual-Marqui et al, 1995) and later called ‘strength’ or Besag factor (Besag, 1986), had minimal effects on smoothing in our dataset. This can be explained by the relative magnitude of the terms in Equation 8. The smoothing algorithm can be read as a Bayesian optimization problem where λ is the log-odds gain, *N*_*b*,κ,*t*_ is the log-prior, and the preceding sum is the negative log-likelihood or residual term. In our case, the log-prior counts the number of microstate labels identical to the current label at time point *t*. Given the magnitude of the residual terms in our data, at λ = 5, a single microstate label identical to *X*_*t*_ within the smoothing window raises the odds of accepting *X*_*t*_ to approximately 150:1. Increasing λ further has negligible effects. Measurable effects would be observed for λ < 1 which we have not found in the microstate literature.

### Template maps

Figure S1 A shows the group level microstate maps from *n* = 201 subjects, obtained from modified K-means clustering. The maps are labelled A-E, according to the maps reported in the literature. Microstate classes are also colour-coded, as indicated by the coloured rectangle above each map. Backfitting and sequence smoothing are visualized in Figure S1 B-D, where the microstate classes are presented as coloured areas under the GFP timecourse. The GFP timecourse is identical in B-D. Panel B shows a non-smoothed sequence, where every temporal sample has been fitted. Panel C shows the results after parametric smoothing of the sequence in B with *b* = 5 samples window half-size (window size 44 ms), and panel D shows the results of interpolation between GFP peaks. It is evident that parametric smoothing and interpolation increase microstate duration and reduce occurrence rates, compared to the non-smoothed sequence in B.

**Fig. S1.**
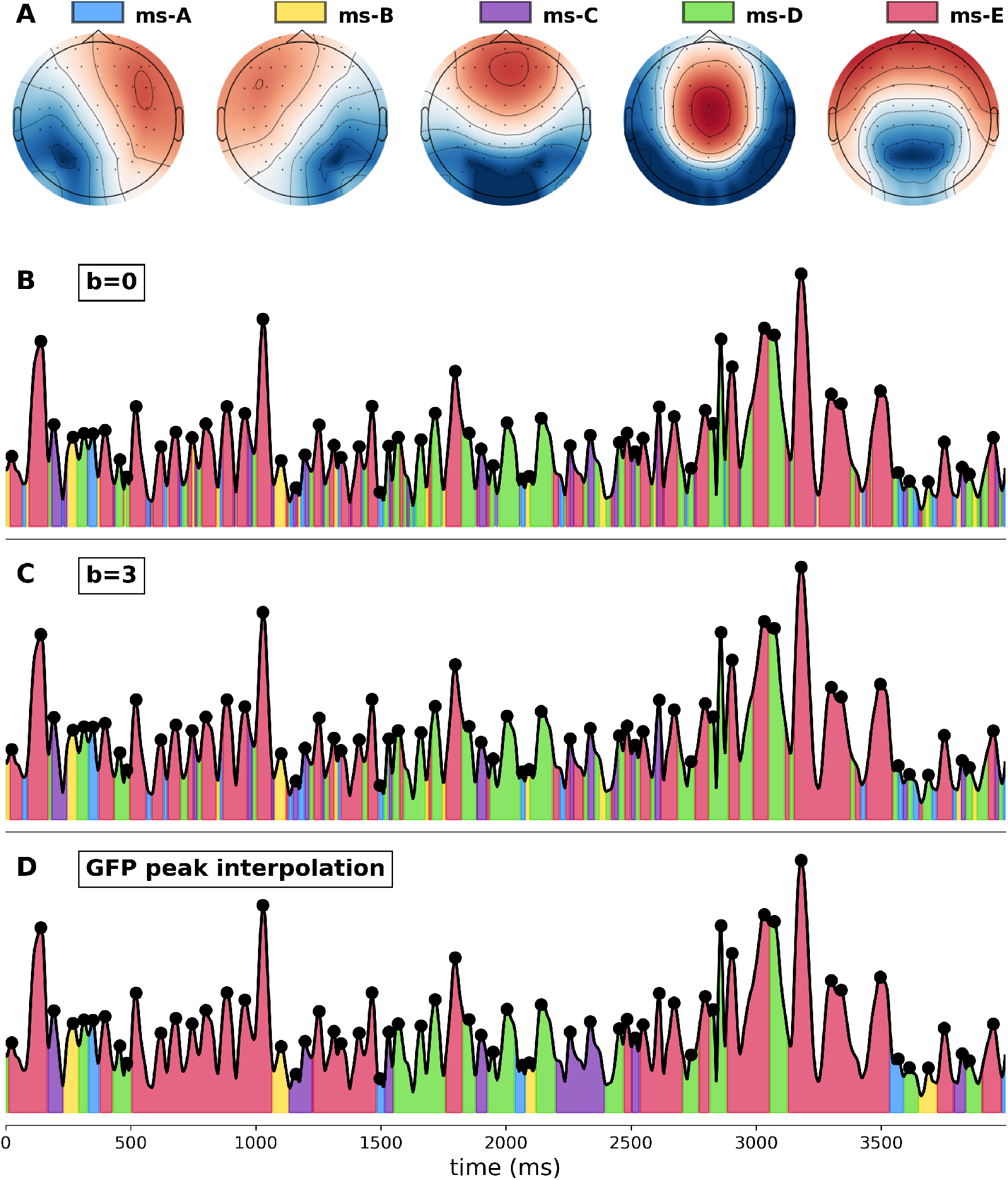
**A**: Group microstate maps (*N* = 201) for *K* = 5 microstate classes, labelled as ms-A to ms-E. The color bars above the maps are used to identify these classes in panels **B** - **D.**Orientation: top=frontal, bottom=occipital, left/right as shown. Panels **B**-**D** show the same GFP time course example (black) and the corresponding microstate class using the color scheme shown in **A. B**: Microstate sequence without smoothing (*b* = 0) or interpolation. **C**: Microstate sequence after parametric smoothing using window half-size *b* = 5 samples, or window size 44 ms. **D**: GFP-peak interpolated microstate sequence.

## References

Al Zoubi O, Mayeli A, Tsuchiyagaito A, et al (2019) Eeg microstates temporal dynamics differentiate individuals with mood and anxiety disorders from healthy subjects. Frontiers in Human Neuroscience 13:56. 10.3389/fnhum.2019.00056

Babayan A, Erbey M, Kumral D, et al (2019) A mind-brain-body dataset of mri, eeg, cognition, emotion, and peripheral physiology in young and old adults. Scientific Data 6:180308. 10.1038/sdata.2018.308

Besag J (1986) On the statistical analysis of dirty pictures. Journal of the Royal Statistical Society Series B (Methodological) 48(3):259–302

Bodda S, Diwakar S (2025) Interconnections and global transitions among functional states encode activity-related dynamics as brain topology changes after yoga training. Scientific Reports 15:16845. 10.1038/s41598-025-00134-y

Bréchet L, Michel CM (2022) Eeg microstates in altered states of consciousness. Frontiers in Psychology 13. 10.3389/fpsyg.2022.856697, URL http://dx.doi.org/10.3389/fpsyg.2022.856697

Cai Y, Huang D, Chen Y, et al (2018) Deviant dynamics of resting state electroencephalogram microstate in patients with subjective tinnitus. Frontiers in Behavioral Neuroscience 12:122. 10.3389/fnbeh.2018.00122

Cai Y, Chen S, Chen Y, et al (2019) Altered resting-state eeg microstate in idiopathic sudden sensorineural hearing loss patients with tinnitus. Frontiers in Neuroscience 13:443. 10.3389/fnins.2019.00443

Chivu A, Pascal SA, Damborsk A, et al (2024) Eeg microstates in mood and anxiety disorders: A meta-analysis. Brain Topography 37:357–368. 10.1007/s10548-023-00999-0

Coquelet N, De Tige X, Roshchupkina L, et al (2022) Microstates and power envelope hidden markov modeling probe bursting brain activity at different timescales. NeuroImage 247:118850. 10.1016/j.neuroimage.2021.118850

da Cruz JR, Favrod O, Roinishvili M, et al (2020) Eeg microstates are a candidate endophenotype for schizophrenia. Nature Communications 11:3089. 10.1038/s41467-020-16914-1

Damborská A, Piguet C, Aubry JM, et al (2019) Altered electroencephalographic resting-state large-scale brain network dynamics in euthymic bipolar disorder patients. Frontiers in Psychiatry 10:826. 10.3389/fpsyt.2019.00826

D’Croz-Baron DF, Baker M, Michel CM, et al (2019) Eeg microstates analysis in young adults with autism spectrum disorder during resting-state. Frontiers in Human Neuroscience 13:173. 10.3389/fnhum.2019.00173

Férat V, Scheltienne M, Brunet D, et al (2022a) Pycrostates: a Python library to study EEG microstates. Journal of Open Source Software 7(78):4564. 10.21105/joss.04564, URL http://dx.doi.org/10.21105/joss.04564

Férat V, Seeber M, Michel CM, et al (2022b) Beyond broadband: Towards a spectral decomposition of electroencephalography microstates. Human Brain Mapping 43:3047–3061. 10.1002/hbm.25834

de la Fuente LA, Zamberlan F, Bocaccio H, et al (2023) Temporal irreversibility of neural dynamics as a signature of consciousness. Cerebral Cortex 33:1856–1865. 10.1093/cercor/bhac177

Gao F, Jia H, Feng Y (2018) Microstate and omega complexity analyses of the resting-state electroencephalography. Journal of Visualized Experiments : JoVE 10.3791/56452

González-Villar AJ, Trianes Y, Gómez-Perretta C, et al (2020) Patients with fibromyalgia show increased beta connectivity across distant networks and microstates alterations in resting-state electroencephalogram. NeuroImage 223:117266. 10.1016/j.neuroimage.2020.117266, URL https://www.sciencedirect.com/science/article/pii/S1053811920307527

Gramfort A (2013) MEG and EEG data analysis with MNE-Python. Frontiers in Neuroscience 7. 10.3389/fnins.2013.00267, URL http://dx.doi.org/10.3389/fnins.2013.00267

Hao Z, Zhai X, Cheng D, et al (2022) Eeg microstate-specific functional connectivity and stroke-related alterations in brain dynamics. Frontiers in Neuroscience 16:848737. 10.3389/fnins.2022.848737

Hermann G, Tdt I, Tagliazucchi E, et al (2024) Propofol reversibly attenuates short-range microstate ordering and 20 hz microstate oscillations. Brain Topography 37(2):329–342. 10.1007/s10548-023-01023-1, URL http://dx.doi.org/10.1007/s10548-023-01023-1

Jia H, Yu D (2019) Aberrant intrinsic brain activity in patients with autism spectrum disorder: Insights from eeg microstates. Brain Topography 32:295–303. 10.1007/s10548-018-0685-0

Jia W, von Wegner F, Zhao M, et al (2021) Network oscillations imply the highest cognitive workload and lowest cognitive control during idea generation in open-ended creation tasks. Scientific Reports 11:24277. 10.1038/s41598-021-03577-1

Koenig T, Lehmann D, Merlo MC, et al (1999) A deviant EEG brain microstate in acute, neuroleptic-naive schizophrenics at rest. Eur Arch Psychiatry Clin Neurosci 249(4):205–211

Koenig T, Prichep L, Lehmann D, et al (2002) Millisecond by millisecond, year by year: normative EEG microstates and developmental stages. NeuroImage 16(1):41–48

Krylova M, Alizadeh S, Izyurov I, et al (2021) Evidence for modulation of eeg microstate sequence by vigilance level. NeuroImage 224:117393. 10.1016/j.neuroimage.2020.117393

Lehmann D, Faber PL, Galderisi S, et al (2005) EEG microstate duration and syntax in acute, medication-na ï ve, first-episode schizophrenia: a multi-center study. Psychiatry Research: Neuroimaging 138:141–156

Li Z, Ren G, Liu C, et al (2021) Dysfunctional brain dynamics of parkinson’s disease and the effect of acute deep brain stimulation. Frontiers in Neuroscience 15:697909. 10.3389/fnins.2021.697909

Lian H, Li Y, Li Y (2021) Altered eeg microstate dynamics in mild cognitive impairment and alzheimer’s disease. Clinical Neurophysiology : official journal of the International Federation of Clinical Neurophysiology 132:2861–2869. 10.1016/j.clinph.2021.08.015

Liu Z, Si L, Xu W, et al (2022) Characteristics of eeg microstate sequences during propofol-induced alterations of brain consciousness states. IEEE Transactions on Neural Systems and Rehabilitation Engineering : a publication of the IEEE Engineering in Medicine and Biology Society 30:1631–1641. 10.1109/TNSRE.2022.3182705

Lynn CW, Cornblath EJ, Papadopoulos L, et al (2021) Broken detailed balance and entropy production in the human brain. Proceedings of the National Academy of Sciences of the United States of America 118. 10.1073/pnas.2109889118

Lyons RG (2011) Understanding Digital Signal Processing (3rd Edition), 3rd edn. Prentice Hall

Manasova D, Perl YS, Bruno NM, et al (2025) Dynamics of eeg microstates change across the spectrum of disorders of consciousness. Brain Topography 38:65. 10.1007/s10548-025-01142-x

Michel CM, Bréchet L (2026) Eeg microstates: from methodological foundations to clinical translation. Trends in Neurosciences 49:433–449. 10.1016/j.tins.2026.04.003

Michel CM, Koenig T (2018) EEG microstates as a tool for studying the temporal dynamics of whole-brain neuronal networks: a review. NeuroImage 180:577–593

Michel CM, Brechet L, Schiller B, et al (2024) Current state of EEG/ERP microstate research. Brain Topography 37:169–180. 10.1007/s10548-024-01037-3

Mikutta CA, Knight RT, Sammler D, et al (2024) Electrocorticographic activation patterns of electroencephalographic microstates. Brain Topography 37:287–295. 10.1007/s10548-023-00952-1

Milz P, Faber PL, Lehmann D, et al (2015) The functional significance of EEG microstates-associations with modalities of thinking. NeuroImage 125:643–656

Milz P, Pascual-Marqui RD, Achermann P, et al (2017) The EEG microstate topography is predominantly determined by intracortical sources in the alpha band. NeuroImage 162:353–361

Musaeus CS, Nielsen MS, Høgh P (2019a) Microstates as disease and progression markers in patients with mild cognitive impairment. Front Neurosci 13:563

Musaeus CS, Salem LC, Kjaer TW, et al (2019b) Microstate changes associated with alzheimer’s disease in persons with down syndrome. Frontiers in Neuroscience 13:1251. 10.3389/fnins.2019.01251

Musaeus CS, Engedal K, Hgh P, et al (2020) Changes in the left temporal microstate are a sign of cognitive decline in patients with alzheimer’s disease. Brain and Behavior 10:e01630. 10.1002/brb3.1630

Nagabhushan Kalburgi S, Kleinert T, Aryan D, et al (2023) Microstatelab: The eeglab toolbox for resting-state microstate analysis. Brain Topography 37(4):621–645. 10.1007/s10548-023-01003-5, URL http://dx.doi.org/10.1007/s10548-023-01003-5

Pan DN, Hoid D, Gu RL, et al (2020) Emotional working memory training reduces rumination and alters the eeg microstate in anxious individuals. NeuroImage Clinical 28:102488. 10.1016/j.nicl.2020.102488

Pascual-Marqui RD, Michel CM, Lehmann D (1995) Segmentation of brain electrical activity into microstates: model estimation and validation. IEEE Trans Biomed Eng 42(7):658–665

Perrottelli A, Marzocchi FF, Di Lorenzo G, et al (2025) Detecting electrophysiological alterations in psychiatric disorders through event-related microstates: a systematic review. Frontiers in Psychiatry 16:1591079. 10.3389/fpsyt.2025.1591079

Poulsen AT, Pedroni A, Langer N, et al (2018) Microstate eeglab toolbox: An introductory guide. bioRxiv 10.1101/289850, URL https://www.biorxiv.org/content/early/2018/03/27/289850, https://www.biorxiv.org/content/early/2018/03/27/289850.full.pdf

Roldán E, Parrondo JM (2010) Estimating dissipation from single stationary trajectories. Physical Review Letters 105(15):150607. 10.1103/PhysRevLett.105.150607, URL https://link.aps.org/doi/10.1103/PhysRevLett.105.150607

Schumacher J, Peraza LR, Firbank M, et al (2019) Dysfunctional brain dynamics and their origin in Lewy body dementia. Brain 142(6):1767–1782

Shi W, Chen F, Cao Y, et al (2026) Research on the syntax of eeg microstates: A comprehensive analysis of epilepsy and psychosis of epilepsy. Brain Research Bulletin 234:111720. 10.1016/j.brainresbull.2026.111720

Sleigh J, Hight D (2021) Is complexity complicated? British Journal of Anaesthesia 127:173–174. 10.1016/j.bja.2021.05.014

Tarailis P, Koenig T, Michel CM, et al (2023) The functional aspects of resting eeg microstates: A systematic review. Brain Topography 10.1007/s10548-023-00958-9

Van de Ville D, Britz J, Michel CM (2010) EEG microstate sequences in healthy humans at rest reveal scale-free dynamics. Proc Natl Acad Sci U S A 107(42):18179–18184

von Wegner F, Laufs H (2018) Information-theoretical analysis of EEG microstate sequences in Python. Front Neuroinform 12:30. 10.3389/fninf.2018.00030

von Wegner F, Tagliazucchi E, Brodbeck V, et al (2016) Analytical and empirical fluctuation functions of the EEG microstate random walk - short-range vs. long-range correlations. NeuroImage 141:442–451

von Wegner F, Tagliazucchi E, Laufs H (2017) Information-theoretical analysis of resting state EEG microstate sequences - non-Markovianity, non-stationarity and periodicities. NeuroImage 158:99–111

von Wegner F, Wiemers M, Hermann G, et al (2023) Complexity measures for eeg microstate sequences: Concepts and algorithms. Brain Topography 37(2):296–311. 10.1007/s10548-023-01006-2, URL http://dx.doi.org/10.1007/s10548-023-01006-2

von Wegner F, Hermann G, Tödt I, et al (2025) Higher-order eeg microstate syntax and surrogate testing. Computers in Biology and Medicine

von Wegner F, Hermann G, Tdt I, et al (2026) A quantitative comparison of two methods for higher-order eeg microstate syntax analysis. Brain Topography 39. 10.1007/s10548-026-01196-5

Wiemers MC, Laufs H, von Wegner F (2023) Frequency analysis of eeg microstate sequences in wakefulness and nrem sleep. Brain Topography 10.1007/s10548-023-00971-y

Zanesco AP (2024) Normative temporal dynamics of resting eeg microstates. Brain Topography 37:243–264. 10.1007/s10548-023-01004-4

Zhang M, Li Z, Wang L, et al (2021) The resting-state electroencephalogram microstate correlations with empathy and their moderating effect on the relationship between empathy and disgust. Frontiers in Human Neuroscience 15:626507. 10.3389/fnhum.2021.626507

